# Temporal uncertainty in fossil records can distort distributions and ecological niches during periods of climatic instability

**DOI:** 10.64898/2026.03.23.713783

**Authors:** André M. Bellvé, Val J. P. Syverson, Jessica L. Blois, Marta A. Jarzyna

## Abstract

Reliable models of species niches and distributions depend on accurately matching occurrences to environments via spatial and temporal coordinates. For fossil occurrences, time-averaging and age uncertainty can create mismatches between fossils and their associated environments, distorting inferred niches and distributions. Using a virtual ecology approach, we assessed how temporal uncertainty (±200 years to the full late Quaternary) influences niche and distribution estimates for four virtual species centered on three periods: Holocene (6,000 y.b.p), deglacial (13,500 y.b.p.), and Last Glacial Maximum (18,000 k.y.b.p.). We compared ‘uncertain’ estimates, derived by matching occurrences with environmental layers drawn from different times within each uncertainty window, against ‘true’ niches and distributions. We found that during environmentally stable intervals, niches and distributions were robust to temporal uncertainty until it reached ±2500 years. Higher environmental variability reduced accuracy, with the greatest mismatch occurring during the deglacial. These results demonstrate both the promise and limitations of paleodistribution reconstruction.

## Introduction

Paleo-archives (e.g., fossils) are an invaluable resource for addressing fundamental ecological and evolutionary questions, offering critical evidence for the abiotic and biotic drivers of extinction and speciation (Grether *et al*. 2024; O’keefe *et al*. 2023; Payne *et al*. 2024) and responses of species and ecosystems to past climatic events (Fordham *et al*. 2020; Meersch *et al*. 2025). For these reasons, the use of the fossil record in ecological niche models (ENMs) has expanded considerably over the past two decades (Blois *et al*. 2025). Incorporating fossils into ENMs offers insights into how species’ niches and distributions have changed over millennia (Fordham *et al*. 2020), often as a result of climatic and human pressures (Bellvé *et al*. 2025b; Canteri *et al*. 2022; Fordham *et al*. 2022; Myers *et al*. 2015). For instance, including fossil and historic occurrences in an ENM—alongside or in place of present-day records—can broaden estimates of abiotic niches, with important implications for species conservation (Bellvé *et al*. 2025b; Lima-Ribeiro *et al*. 2017; Raiter & Hawlena 2024). Despite its utility, the fossil record is subject to unique spatial and temporal biases that can compromise the reliability of ENMs (Barr & Wood 2024; Bellvé *et al*. 2025b, a; Inman *et al*. 2018; Maguire *et al*. 2015), which depend on accurately linking occurrences to environmental conditions through spatial and temporal coordinates.

Fossil occurrences often carry substantial uncertainty in space and time. For instance, preservation potential—the probability that environments allow fossil formation and long-term persistence—varies through time and space, affecting where fossils are preserved and ultimately discovered (Behrensmeyer *et al*. 2000; Darroch *et al*. 2021; Maxwell *et al*. 2018). When ENMs are trained on the fossil record, such preservation biases can distort reconstructions of species’ niches and predictions of their distributions, as environments that are more or less likely to preserve fossils are systematically over- or under-represented (Bellvé *et al*. 2025a). Uncertainty around the dates of fossils may introduce additional distortions: while the spatial coordinates of a fossil occurrence remain fixed, the environment of that location changes through time. Thus, temporal uncertainty implies that the assumed environmental data associated with an occurrence may not accurately reflect the conditions at the time of occurrence (Davis *et al*. 2014; Maguire *et al*. 2015; Myers *et al*. 2015; Reside *et al*. 2010).

Temporal uncertainties in fossil ages arise from many sources, including the precision of radiometric dating techniques (Herrando-Pérez & Stafford Jr. 2025; Wright 2017), the calibration of dates (Reimer *et al*. 2020) and development of a chronology (Parnell *et al*. 2011), or the assignment of ages based on time-averaged stratigraphic units (Behrensmeyer & Chapman 1993; Graham 1993). These uncertainties typically range from centuries to millennia in the ‘near’ past (i.e., the last 130,000 years) but can span millions of years in ‘deep time’, when more precise techniques such as radiocarbon dating are not viable (Blois *et al*. 2025; Walker 2013).

Such temporal uncertainty can distort our understanding of ecological and evolutionary processes. For instance, Tomašových *et al*. (2024) showed that time-averaging of fossil assemblages can alter abundance-diversity relationships, warping our perception of biodiversity across time. Similarly, phylogenetic reconstructions depend on accurate fossil ages to infer the drivers of speciation and extinction (Barido-Sottani *et al*. 2020), as misdated fossils may misalign evolutionary events with relevant biotic or abiotic factors. While the effects of temporal uncertainty are relatively well documented in phylogenetics (e.g., Dos Reis *et al*. 2015; Pol & Norell 2006), their impact on ENMs—which rely on precise correspondence between occurrences and environmental conditions—remains largely unexplored. There are reasons to expect that these impacts could be substantial. For example, during the late Quaternary, the mean annual temperature at a site in North America [1 km × 1 km; 100° W, 40° N] (Karger *et al*. 2023) ranged from 7.9 C° at 12,000 years before present (y.b.p.; where present is 1950 A.D.) to 2.4 C° at 15,000 y.b.p., representing dramatically different environmental conditions within a geologically short 3,000-year time period. As a result, uncertainty in the age estimate of a fossil occurrence could lead to markedly different ENM predictions depending on which specific age is assumed to be ‘true’. Without explicitly accounting for these uncertainties, we risk generating erroneous inferences that could distort our understanding of ecological and evolutionary processes and misinform conservation actions (Johnson *et al*. 2017, 2024).

Here, we investigate how temporal uncertainty in the fossil record impacts our ability to reconstruct niches and predict distributions of four virtual small mammal species in North America, focusing on the last ∼22,000 years of the late Quaternary. Using a virtual ecology approach (Zurell *et al*. 2010), we first created ‘true’ niches and distributions for each virtual species and then compared these with niches and distributions reconstructed from virtual species sampled with varying levels of temporal uncertainty (Fig. 1). We focused on three climatically different time points during the late Quaternary (Clark *et al*. 2012): the relatively warm and stable Holocene interglacial, centered at 6,000 y.b.p.; the climatically variable deglacial transition, centered at 13,500 y.b.p.; and the cold and moderately variable Last Glacial Maximum, centered at 18,000 y.b.p. We expected increasing temporal uncertainty to manifest in three primary ways: 1) increasing distortion of species niche reconstructions, 2) higher distribution prediction errors, and 3) greater impacts on both niche reconstruction and prediction errors under conditions of greater environmental variability.

**Figure 1.**
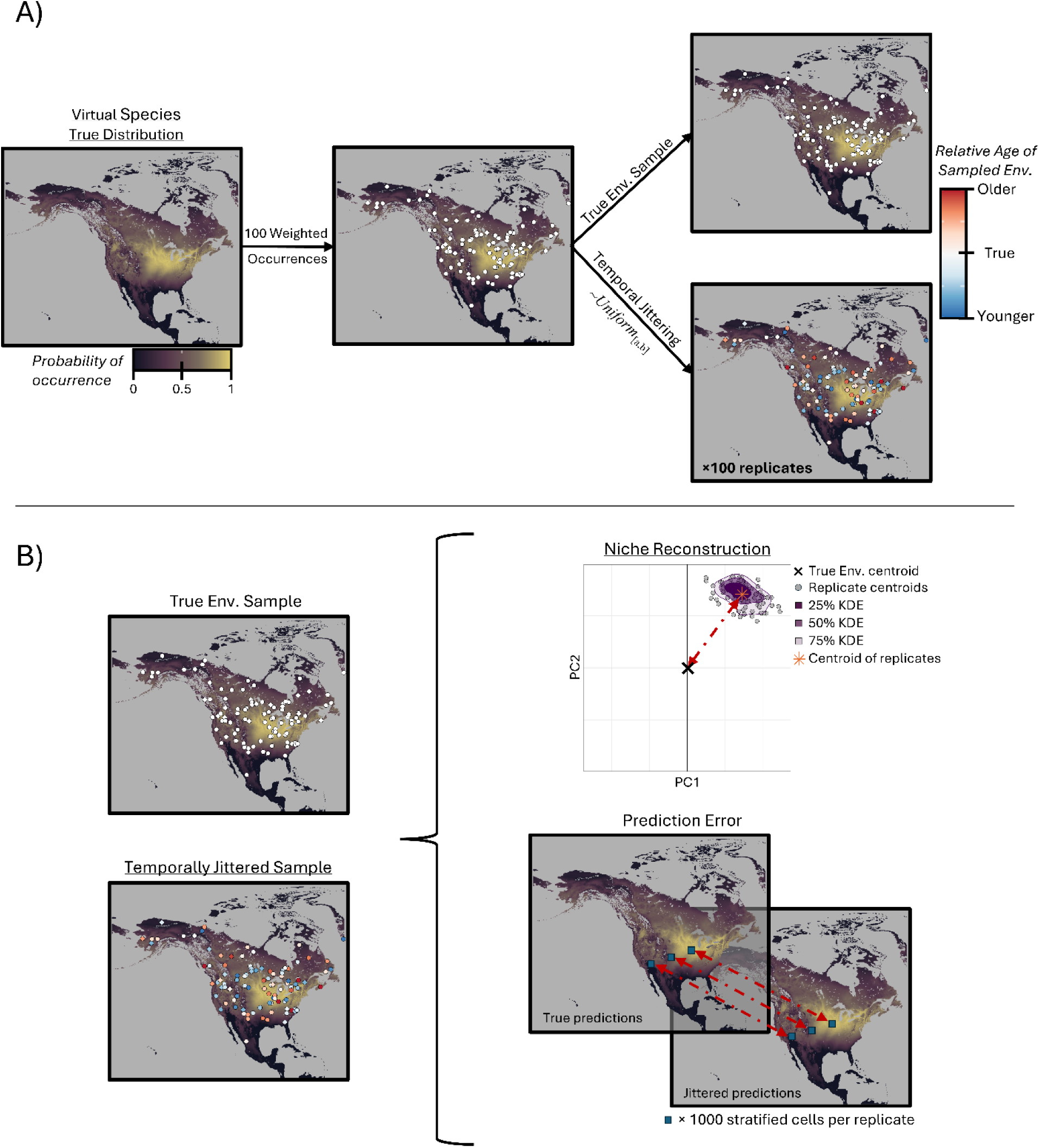
Conceptual workflow of study design. (A) True virtual species distributions were sampled for 100 occurrences, and then temporally jittered by drawing from the uncertainty range using a uniform distribution – a process which was repeated 100 times for each scenario. (B) Models trained on temporally jittered samples were compared to models trained on original true sample using Principal Component Analysis and the distribution predictions from the Maximum Entropy algorithm.

## Methods

### Environmental data

We constructed ENMs for four virtual species of small mammals (< 2 kg adult body mass) using six environmental variables: elevation, terrain slope, mean annual temperature, temperature seasonality, annual precipitation, and precipitation seasonality (see SI.1; Table S1). Together, these variables capture key physical and climatic dimensions of small mammal niches (Beaumont *et al*. 2016) across our study area of North America. All layers were sourced from CHELSA-TraCE21K (Karger *et al*. 2023), except for terrain slope, which we derived from the CHELSA-TraCE21K digital elevation model based on the surrounding eight grid cells. All layers had a 1 km × 1 km spatial resolution and were available at 100-year increments for the last 22,000 years. To mask major water bodies, we used the NaturalEarth ‘North American historic lakes’ shapefile (v5.0.0; naturalearthdata.com) for the present and the Holocene (11,700 y.b.p. - present), assuming relative stability of water bodies over the last 11,700 years. For the deglacial (14,900 – 11,800 y.b.p) and the Last Glacial Maximum (LGM; 22,000 – 15,000 y.b.p.), we additionally masked pluvial water bodies, which were a defining landscape feature during these time periods (v4.0.0; naturalearthdata.com). Glaciers were masked using each time period’s contemporaneous glacier elevation layers from CHELSA-TraCE21K.

### Virtual species creation

Our overarching goal was to assess how temporal uncertainty in fossil dates affects our ability to reconstruct species’ ecological niches and distributions. While it would be ideal to use the actual small mammal fossil record to address this question, range or niche changes in the fossil record could be difficult to disentangle from the effects of temporal uncertainty. To circumvent this limitation, we created four virtual species, modelled on real-world species to ensure ecological realism. We followed the two-step process outlined in Bellvé *et al*. (2025a), using four of the same virtual species. Briefly, we first created ENMs for real contemporary small mammal species, then abstracted them to generate virtual analogs. For the first step of this workflow, we selected four North American small mammal species (SI.2; Fig. S1) from the Global Biodiversity Information Facility (GBIF) database, chosen based on the availability of modern occurrence data, coverage across North America and present-day environments, and representation of diverse ecological habits (Fig. S2). The selected species included two Rodentia (*Microtus pennsylvanicus*, *Neotoma albigula*), one Lagomorpha (*Lepus californicus*), and one Eulipotyphla (*Blarina carolinensis*). Occurrences for each species were used to train MaxEnt models, which were fitted using a five-fold cross-validation (CV). Models were evaluated using the area under the receiver-operator curve (AUC), which ranges between 0.5 (performing no better than chance) and 1 (perfect prediction; Fielding & Bell 1997) and combines measures of sensitivity and specificity. Variable importance was calculated using jackknife CV within each of the models’ five-fold CV replicates (SI.2; Table S3).

We then used the response curves generated from these real species ENMs to create virtual representations of each species’ niches (*virtualspecies* v1.5.1; Leroy *et al*. 2016). Statistical moments derived from these response curves were used to parameterize mathematical functions that approximated each virtual species’ response to environmental variables. These functions were then used to generate probability of occurrence surfaces for each virtual species × time period combination. To assess the performance of these mathematical abstractions, we visually compared the resulting virtual species distributions with those of the real species distributions to ensure similarity. The final output consisted of probability of occurrence layers for each virtual species at three time period midpoints (Holocene – 6,000 y.b.p.; deglacial – 13,500 y.b.p., LGM–18,000 y.b.p.). To denote species’ virtual counterparts, the prefix ‘*V_*’ was added to each species name (e.g., *V_N. albigula*).

### Introducing temporal uncertainty into ecological niche modelling

From each virtual species’ distribution in each time period, we sampled 100 occurrences weighted by species’ probability of occurrence, and 10,000 randomly drawn background points (Barbet-Massin *et al*. 2012). We then trained the ‘true’ ENM (both Maximum Entropy [MaxEnt] and Principal Component Analysis [PCA]) for each virtual species × time period using environmental values drawn from the contemporaneous layers (i.e., the environmental layers corresponding to the same time period as the virtual species occurrences; Fig. 1.A). Next, we created a suite of analogous jittered ENMs, where we introduced temporal uncertainty by randomly reassigning dates to each occurrence from uniform distributions over specified uncertainty ranges (± 200, 400, 600, 800, 1000, 1500, 2500, 4000 years, and the entirety of the late Quaternary [LQ; ∼ the last 22,000 years)], centered on each time period, in 100 year increments. We refer to this process of introducing temporal uncertainty as ‘jittering’ throughout the manuscript. Environmental values were then extracted from the 100 jittered occurrences for each time period (Fig. 1.A). In essence, spatial coordinates remained fixed for the set of occurrences while temporal uncertainty was added to the associated environmental samples. For each combination of virtual species, time period, and temporal uncertainty range, we replicated the entire temporal jittering procedure (i.e., adding temporal uncertainty to each occurrence in the original sample) 100 times, to measure the variability with which these random uniform draws influence our results. We note that our study design is the inverse of reality; in the real world, fossils are deposited through time, but ENM studies often treat fossils as falling within a single time bin (Blois *et al*. 2025) (e.g., the Holocene), ignoring the possibility that ‘true’ environmental conditions may not align with the chosen midpoint of a time bin. While our framework reverses this assumption, the resulting insights into distributions and niches are directly transferable. We measured the effects of temporal uncertainty on ENMs by comparing our ‘true’ and temporally jittered models in two primary ways: niche reconstruction accuracy and distribution prediction error (Fig. 1.B).

### Niche reconstruction accuracy

To characterize each virtual species’ ‘true’ environmental niche, we centered and scaled the environmental values of each time period, and fitted them with a covariance-based Principal Component Analysis (PCA; hereafter, ‘true PCA’). Each of these true virtual species PCAs was paired with the temporally jittered PCA replicate for each temporal uncertainty range. We superimposed the 100 temporally jittered occurrences of each replicate onto its respective true PCA space and then calculated the replicate centroid. Finally, we calculated the distance from the center of these replicate centroids to the true centroid, as a measure of niche reconstruction accuracy.

### Distribution prediction error

We created models using the maximum entropy algorithm for both the true occurrences (the ‘true MaxEnt’ model) and the temporally jittered occurrences (the ‘temporally jittered MaxEnt’ models). For the true MaxEnt model (i.e., the model based on the environmental values without temporal uncertainty), we randomly selected 100 cells from the predicted distribution at 0.01 probability of occurrence at increments between 0 – 1. This approach reduced the computational burden of predicting the value for the ∼27.5 million non-NA cells in our rasters for each of the 10,800 sample models (3 time periods × 4 virtual species × 9 uncertainty ranges × 100 replicates), while maintaining a representative set of environmental conditions across the virtual species’ distributions. We then predicted the probability of occurrence for each of these randomly selected cells using the temporally jittered MaxEnt models and compared these predictions to the true probability of occurrence. For each model replicate, distribution prediction errors were summarized by deriving their root mean squared error (RMSE) for each model replicate.

### Environmental variability

To quantify environmental variability, we sampled 1,000 cells from each virtual species’ true distribution weighted by the probability of occurrence and extracted the ‘true’ value of all environmental layers for the time period’s midpoint. For each sampled cell, we also extracted its environmental values from all environmental layers within the temporal uncertainty range and calculated their difference from the true values. This process resulted in a set of differences for each environmental variable, corresponding to the number of time periods within the temporal uncertainty range (e.g., for the temporal uncertainty range ± 200, there were four total differences: focal time – 100 years; focal time – 200 years; focal time + 100 years; focal time + 200 years). We then calculated the quartiles of these differences across all cells in the sample and computed the interquartile range (IQR; 0.25-0.75). The IQRs were normalized to a 0-1 scale for each environmental variable and species using feature scaling (Eq. 1), so that environmental variables were commensurate with each other.

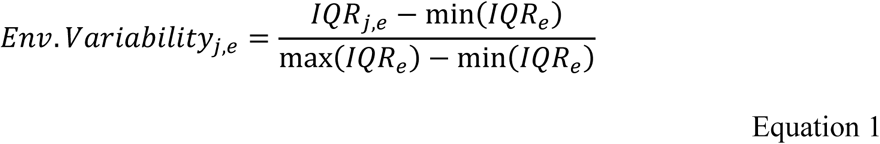

Where *j* indicates the *j*^th^ sampled cell, and *e is* a given environmental variable. All scaled environmental variability values were then summed to yield a single composite metric of environmental variability for a temporal uncertainty range, and we repeated the procedure for each level of temporal uncertainty. Note that we use the phrase “environmental variability” throughout the manuscript as a shorthand to refer to the scaled range in environmental differences from the time period’s midpoint. Thus, the calculated environmental variability over the full LQ period differs among the LGM, deglacial, and Holocene time periods, because each is measured relative to the environment at its midpoint.

### Implementation

All data manipulation and analyses were performed in R (v4.5.0; R Core Team 2025) using an RStudio interface (v2025.05.1 Build 513 “Mariposa Orchid”; Rstudio Team 2025). Large-scale simulations were run in base R in a Linux environment through the Ohio Supercomputer Centre. See SI.3 for more details.

## Results

### Niche reconstruction accuracy

Total variation captured by the first two PCs calibrated on the true occurrence data ranged between 57.5% - 76.6%, across all species and time periods (see SI.4; Fig. S3), indicating that biplots of these two PCs represent species’ environmental niches well. Across all species and time periods, the average distance between the temporally biased niche centroid and the true centroid increased with temporal uncertainty (Fig. 2, Fig. 3), indicating that temporal uncertainty skewed reconstructions of the species niches. However, the magnitude of this effect varied across uncertainty levels and time periods. For example, jittering occurrences across the last 22,000 years (i.e., the entire LQ) resulted in particularly poor reconstructions of Holocene and LGM niches, although the strength of this effect differed among individual species (Fig. 2).

**Figure 2.**
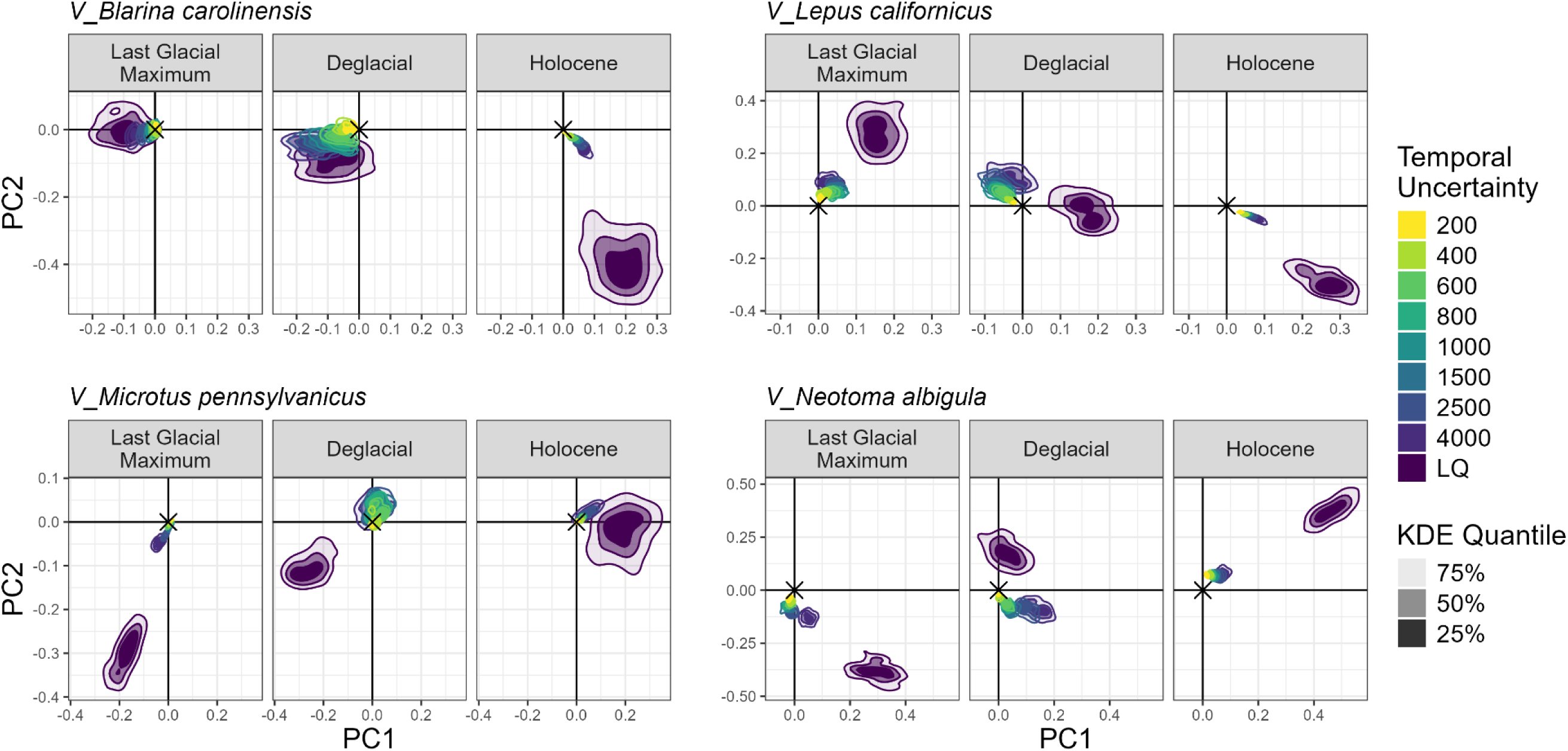
Kernel density estimate bands (25%, 50%, 75%) of the replicate centroids for each level of temporal uncertainty (±), virtual species, and time period. The colour of the KDE bands indicates the level of temporal uncertainty, and their opacity shows the KDE quantiles. The black ‘X’ at the centre of each plot shows the centroid of the true scaled and centred environmental values of the original occurrences, which were used to calibrate the covariance-based Principal Component Analysis.

**Figure 3.**
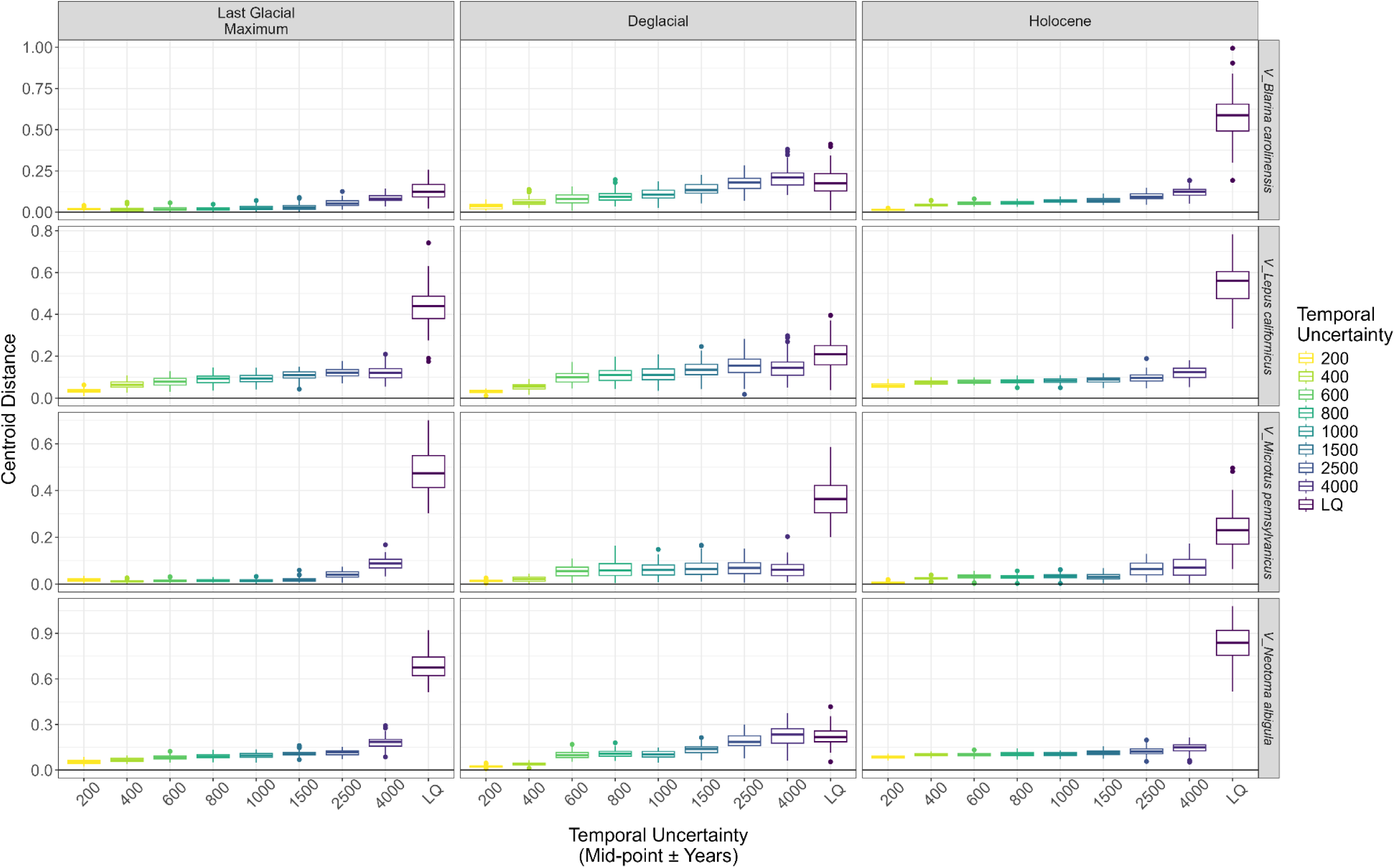
Boxplot of replicate centroid distances between the true and temporally jittered samples for each virtual species and time period replicate, with the colour of the boxes indicating the temporal uncertainty range of the jittered MaxEnt models. Note that the centroid distances are only directly comparable within a species and time period and should not be compared across species or time periods, as each individual PCA was centered on a different set of data.

Temporal uncertainty introduced a consistent, non-random bias during the Holocene and LGM, with jittered niche centroids shifting systematically away from the true niche (Figs. 2, 3). In contrast, the effect of temporal uncertainty during the deglacial period was less consistent, with jittered centroids tending to orbit the true centroid in some cases (Fig. 2); consequently, centroid distance showed a hump-shaped pattern (Fig. 3). This difference between the time periods is likely due to the deglacial period representing an environmental midpoint between the LGM and the Holocene.

### Distribution prediction error

Median distribution prediction error increased with the level of temporal uncertainty (Fig. 4). For the Holocene, median prediction error for all species remained relatively low (≤ ∼ 0.05), until temporal uncertainty reached 2,500 years—except for *V_N. albigula,* which maintained low prediction errors until temporal uncertainty spanned the entire late Quaternary. In contrast, for the deglacial, median prediction error increased steadily with temporal uncertainty across all species and showed relatively steady increases at relatively low temporal uncertainties (i.e., ± 600 years), indicating a stronger effect of temporal uncertainty on prediction accuracy of species’ distributions during this period. During the LGM, the median prediction error increased more gradually with temporal uncertainty than during the deglacial period, and reached the highest errors for all species once temporal uncertainty encompassed the entire late Quaternary (Fig. 4).

**Figure 4.**
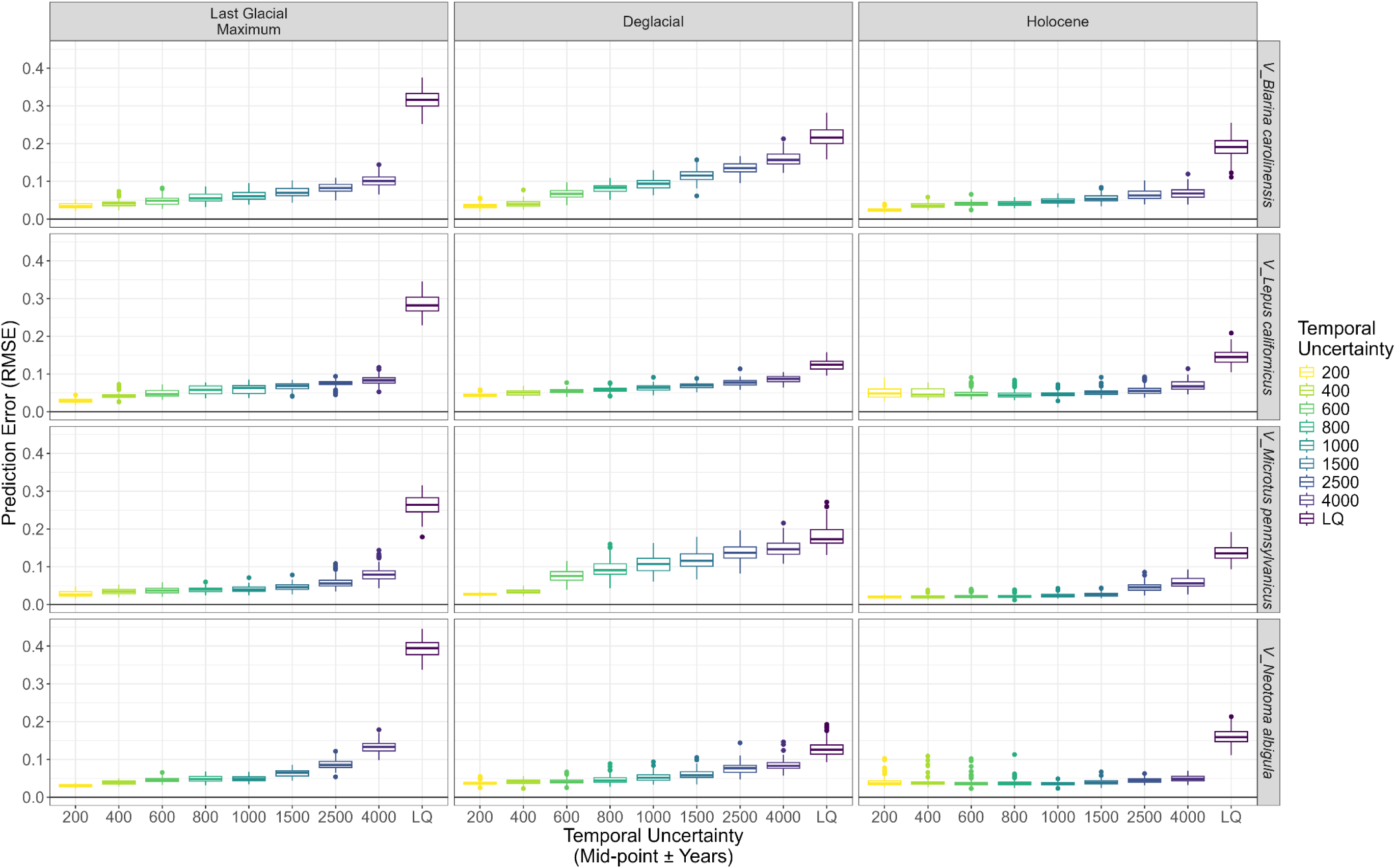
Boxplot of prediction error (root mean square error) of the temporally jittered MaxEnt models from the true MaxEnt model predictions for each virtual species and time period replicate, with the colour of the boxes indicating the temporal uncertainty range of the jittered MaxEnt models.

### Environmental variability

Environmental variability increased monotonically with temporal uncertainty across time periods and species (Fig. 5). Environmental variability was low around the Holocene midpoint, until uncertainty reached ± 2,500 years, when environments for all species showed a step increase in variability (Fig. 5). In contrast, the deglacial period generally showed the greatest rate of increase in environmental variability with temporal uncertainty, especially at low levels of uncertainty.

**Figure 5.**
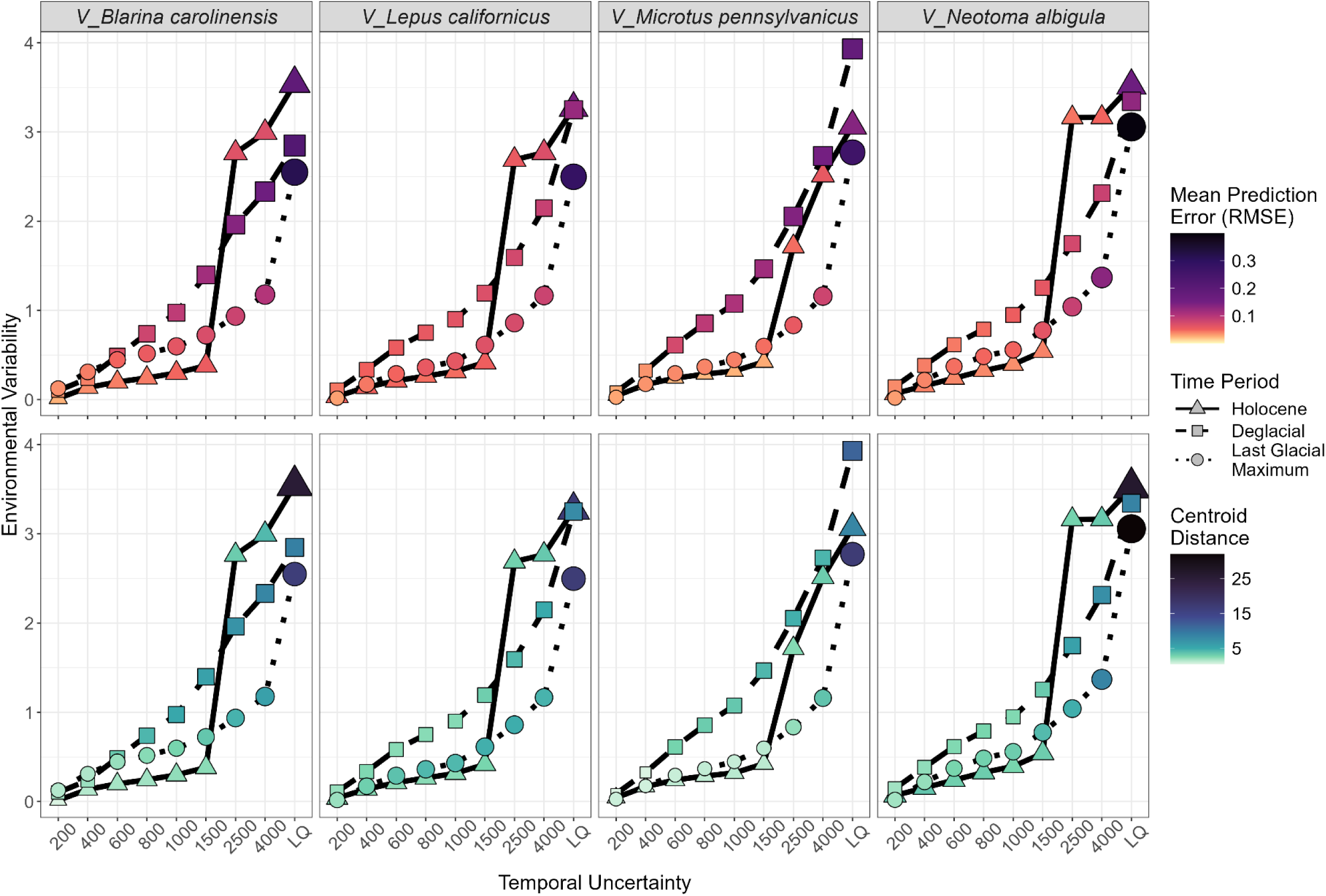
Environmental variability plotted against temporal uncertainty, with point fill colour and size showing average prediction error and centroid distance, respectively. Line type and shape indicate the time period and are facetted by species. Environmental variability was scaled on a species-by-environmental-variable basis, so scaled environmental variability values among species do not necessarily indicate equivalent changes in the original environmental variables.

The LGM exhibited an intermediate pattern, with moderate and steady increases in environmental variability as temporal uncertainty increased. However, environmental variability rose more sharply when temporal uncertainty spanned the entire LQ, likely due to missing data points associated with glacial and sea masking (see SI.5).

Across all species and time periods, mean prediction error and centroid distance increased with environmental variability, though the rate of increase differed among time periods and species. Generally, mean prediction error and mean centroid distance increased more rapidly with temporal uncertainty during the deglacial period (Fig. 5). Exceptions to this were seen in *V_L. californicus* and *V_N. albigula*, where mean prediction error and centroid distance increased more rapidly at low levels of temporal uncertainty during the LGM, irrespective of the higher environmental variability of other periods, although this may partly be the effect of occurrences being lost due to changes in available habitat (see Supplementary Information). In contrast, the Holocene saw very modest increases in mean prediction error and centroid distance across increasing levels of environmental variability, until temporal uncertainty encompassed the entire late Quaternary across all species (Fig. 5).

## Discussion

The fossil record is critical to our understanding of how species respond to environmental change, and to generating robust predictions of future distributions (Blois *et al*. 2025). We demonstrate that reconstructions of species niches and distributions using fossil occurrences are moderately robust to the levels of temporal uncertainty inherent in the fossil record, although this robustness depends on the focal time period and, critically, the extent of environmental variability relative to the temporal uncertainty of fossil ages. For the Holocene and the LGM, temporal uncertainties of less than ± 2,500 years had generally minor effects on species niche reconstructions and predictions of their distributions, suggesting that ENMs are relatively robust to such uncertainties. This level of uncertainty falls well within the typical accuracy of radiocarbon dating for specimens and sites in Holocene, although it is less achievable for the LGM (Syverson *et al*. 2026). In contrast, niche and distribution reconstructions for the deglacial time period showed more variable responses: centroid distance and prediction error increased more rapidly with temporal uncertainty for most species (Figs. 3, 4). We also show that niche reconstruction and distribution predictions were not accurate when temporal uncertainty encompassed the entire late Quaternary (Figs. 2-4). Crucially, our results link the negative effects of temporal uncertainty on niche reconstruction and distribution prediction to the magnitude of environmental variability, demonstrating that the impacts of temporal uncertainty on ENMs are not constant through time (Fig. 5). These findings are the first to underscore the importance of explicitly accounting for temporal uncertainty when assessing niche dynamics or distributional change across periods characterized by high environmental variability.

In reconstructions of fossil niches and distributions, temporal uncertainty is influenced by the temporal information used to assign occurrence ages. Relying on directly dated specimens for a taxon enables more precise alignment between occurrence and environment conditions, whereas the practice of assigning fossil occurrences to broad time bins or biostratigraphic units—and associating them with the corresponding environmental conditions drawn from, e.g., the time period or unit midpoint—introduces greater temporal uncertainty and can lead to less precise niche or distribution inference. As such, studies that have reported apparent changes in niche breadth or shifts in species ranges based on fossil-trained ENMs (e.g., Phelps *et al*. 2020; Rindel *et al*. 2021; Wendt *et al*. 2022) may, in some cases, reflect artifacts arising from inaccurate fossil occurrence ages rather than true ecological change. Likewise, the mixed temporal transferability often observed in models built using fossil occurrences (Maguire *et al*. 2016; Veloz *et al*. 2012; Williams *et al*. 2013) may stem, at least in part, from these distortions, underscoring the importance of explicitly accounting for temporal uncertainty when interpreting past species–environment relationships.

Our study focused on a commonly studied ‘near-time’ period (the late Quaternary; Blois *et al*. 2025) and on virtual fossil occurrences with relatively small temporal uncertainties. Nevertheless, our findings have clear ramifications for research using deep-time fossils, where both fossil age estimates and environmental reconstructions are considerably less certain (Maguire *et al*. 2015). Many deep-time studies rely on fossil-trained ENMs to understand how niche stability or breadth relate to extinction and speciation rates (e.g., Chiarenza *et al*. 2019; Flannery-Sutherland *et al*. 2025; Foffa *et al*. 2025; Parker *et al*. 2018; Stigall 2012; Timmermann *et al*. 2022; Walls & Stigall 2011; Waterson *et al*. 2016). We show, however, that niche reconstructions can vary markedly based solely on the assumed dates of fossil occurrences and their associated environmental conditions, potentially placing such studies at risk of drawing spurious conclusions about evolutionary processes if the environmental niche is not developed appropriately. Our intention is not to critique these previous studies—all of which have provided important insights into deep-time dynamics—but rather to emphasize the need for caution and thoughtful consideration of temporal uncertainty and the magnitude of environmental variability when drawing conclusions from fossil-based models.

The effects of temporal uncertainty on species’ niche centroids and prediction errors are substantially greater than those linked to preservation potential, demonstrated in a related study of the same virtual species and time periods (Bellvé *et al*. 2025a). Although the impacts of preservation potential and temporal uncertainty were assessed independently in these studies, these factors almost certainly interact in reality, given that preservation potential is partially a function of the environment (Bellvé *et al*. 2025a; Noto 2011). Such interactions may further compromise reconstructions of species’ niches and distributions if not explicitly accounted for. While disentangling the joint effects of preservation potential and temporal uncertainty lies beyond the scope of this study, we encourage future research to quantify and integrate these processes more explicitly. We note that while preservation potential can be addressed by weighting background point sampling to represent the sampling biases (Bellvé *et al*. 2025a), addressing temporal uncertainties may be less tractable within the current ENM frameworks, further necessitating research into approaches for explicitly accounting for temporal uncertainty.

Encouragingly, our results suggest that studies using fossils with temporal uncertainties spanning periods of relative environmental stability (e.g., the Holocene) are unlikely to mischaracterize niches or distributions. Nonetheless, we caution against making unwarranted assumptions in either direction and emphasize that apparent niche and range changes inferred from fossil occurrences should be carefully scrutinized and thoroughly validated. At a minimum, fossil occurrences with large temporal uncertainties, or those encompassing periods of marked environmental variability, should be tested to determine whether their inclusion or exclusion alters results and conclusions. Because the fossil record is often sparse for some species and time periods, simply excluding some occurrences may not be feasible. To address this, we recommend conducting sensitivity analyses for niche inference and distribution predictions by varying the dates of high-uncertainty samples within their error bounds or by drawing from probability distributions associated with their chronology, rather than assuming a single date is accurate.

Similarly, extracting environmental predictors using a common temporal midpoint has clear potential to bias inference; instead, the best estimate of each fossil’s age should be used and, where possible, aligned with increasingly fine-resolution paleo-climate reconstructions (e.g., Fordham *et al*. 2022). Sensitivity analyses can also be assessed alongside independent lines of evidence for species niches and distributions—such as phylogenetic data, genomic data, or expert knowledge—and potentially integrated using tools such as Approximate Bayesian Computation to provide more robust insights into the drivers of ecological and evolutionary change (Hoban *et al*. 2019; Naughtin *et al*. 2025; Razgour *et al*. 2013).

Finally, our results can also guide the strategic allocation of resources for fossil age estimation. Constraints on funding, availability of material, and scientific priorities often lead to differential availability of dates across species, times, and sites (Blaauw 2012; Bronk Ramsey 2008; Meltzer & Mead 1985; Saltré *et al*. 2015). We suggest that the effects of temporal uncertainty can help identify which specimens should be prioritized for dating to maximize impact. For instance, fossil occurrences with very large temporal uncertainties—such as those assigned ages based on associated indirect dates or biostratigraphy—or those likely to fall within periods of high environmental variability should be prioritized, as they have the greatest potential to distort ENMs. Ideally, continued advances in dating technologies and chronology techniques will yield increasingly accurate and precise fossil age estimates (e.g., Herrando-Pérez & Stafford 2025; Syverson *et al*. 2026), reducing the influence of temporal uncertainty on niche and distribution reconstructions. For deep-time fossils, technological advances in dating techniques and stratigraphy will also help, but other methods for inferring age, such as estimating fossil ages using Bayesian phylogenetics, may complement these (Drummond & Stadler 2016).

The use of the fossil record in ENMs has grown steadily and is likely to accelerate as paleoecologists and paleobiologists increasingly recognize its value for understanding how species’ niches evolve and respond to environmental change. As their application expands, explicitly accounting for the temporal uncertainty inherent in fossil data will be essential to ensure that ENMs provide robust insights into past species’ niches and distributions. Ultimately, because fossils are central to understanding long-term ecological and evolutionary dynamics (Dietl & Flessa 2011; Fordham *et al*. 2020), there is no path forward without making careful consideration of temporal uncertainty a central aspect of bias correction.

## Authorship Statement

All authors collectively conceptualised the work. AMB designed the virtual ecology experiment, prepared the necessary external data, conducted all analyses, created the figures, and wrote the first draft of the manuscript. MAJ supervised and MAJ and JLB funded the work through a grant from the National Science Foundation (NSF) Division of Earth Sciences (EAR) 2149419 to MAJ and NSF EAR 2149416 to JLB. All authors contributed substantially to the revisions. No generative AI was used in this analysis or preparation of the publication.

## Data Statement

All code necessary to replicate our virtual ecology experiment is available on GitHub (https://github.com/AndreMBellve/temporal_uncertainty). The environmental data necessary for our analysis is available from Karger *et al*. (2023).

## Supplementary Information

### SI.1 – Environmental Data

All environmental layers had a spatial resolution of 1 km × 1 km and were stored in the World Geodetic System 1984 (ESPG: 4326). We restricted our analysis to North America by cropping all layers to a bounding box of xmin = -180, xmax = -50, ymin = 10, ymax = 85.

**Table S1.** Environmental predictors, their respective units and variable ranges for each time period modelled with the periods mid-point.

| Time Period | Environmental Predictor<br>(minimum - maximum) |  |  |  |  |  |
| --- | --- | --- | --- | --- | --- | --- |
|  | Elevation (m) | Slope (°) | Annual Mean<br>Temperature (C°) | Temperature<br>Seasonality (S.D.<br>of monthly means) | Annual<br>Precipitation<br>(kg/m<sup>2</sup>/year) | Precipitation<br>Seasonality<br>(Coefficient of<br>Variation) |
| Present<br>(2000 AD) | -1617 – 5433 | 0 – 53.2 | -21.9 – 31.3 | 25 – 1821 | 0 – 5391 | 5.2 – 134.4 |
| Holocene<br>(6,000 y.b.p.) | -1617 – 5204 | 0 – 56.8 | -22.3 – 31.6 | 27 – 2005 | 0 – 6247 | 6.6 – 163.8 |
| Deglacial<br>(13,500 y.b.p.) | -1617 – 4712 | 0 – 48.6 | -27.5 – 30.6 | 63 – 1911 | 0 – 7150 | 9.9 – 139.2 |
| Last Glacial Maximum<br>(18,000 y.b.p.) | -1617 – 4578 | 0 – 49.9 | -35.0 – 27.6 | 51 – 1865 | 0 – 11378 | 12.0 – 150.0 |

### SI.2 – Virtual species creation

We only retained occurrences that had geographic coordinates, were present on the North American continent, were not fossil specimens, and were recorded between 1950 and 2023. All occurrences were downloaded on 5 October 2023 at 9:30 AM Eastern Standard Time. We further filtered all occurrences by the IUCN expert range maps (https://www.iucnredlist.org/resources/spatial-data-download) to remove any potentially erroneous occurrences; duplicates were also removed (Table S2). We used these real species occurrence data to train MaxEnt ecological niche models, separately for each species. We randomly sampled background points in numbers equal to the number of respective species’ presences. Background points with the same cell ID as presences were excluded. A summary of models performance and predictor importance are shown in Table S3.

**Table S2.**
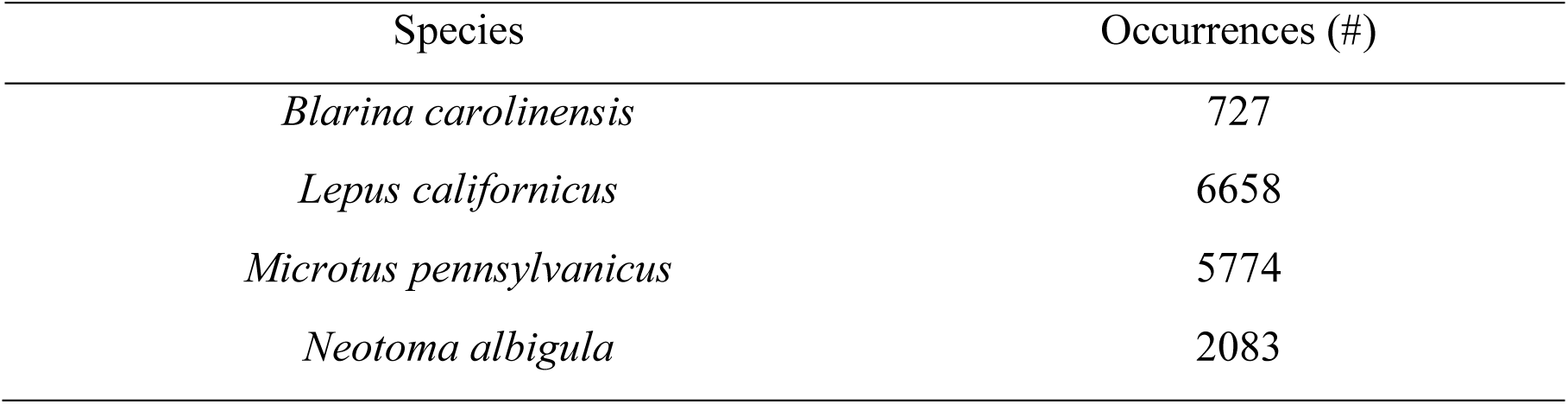
Species we created ENMs for and the number of occurrences retrieved from the GBIF database within the IUCN range maps for North America.

**Figure S1.**
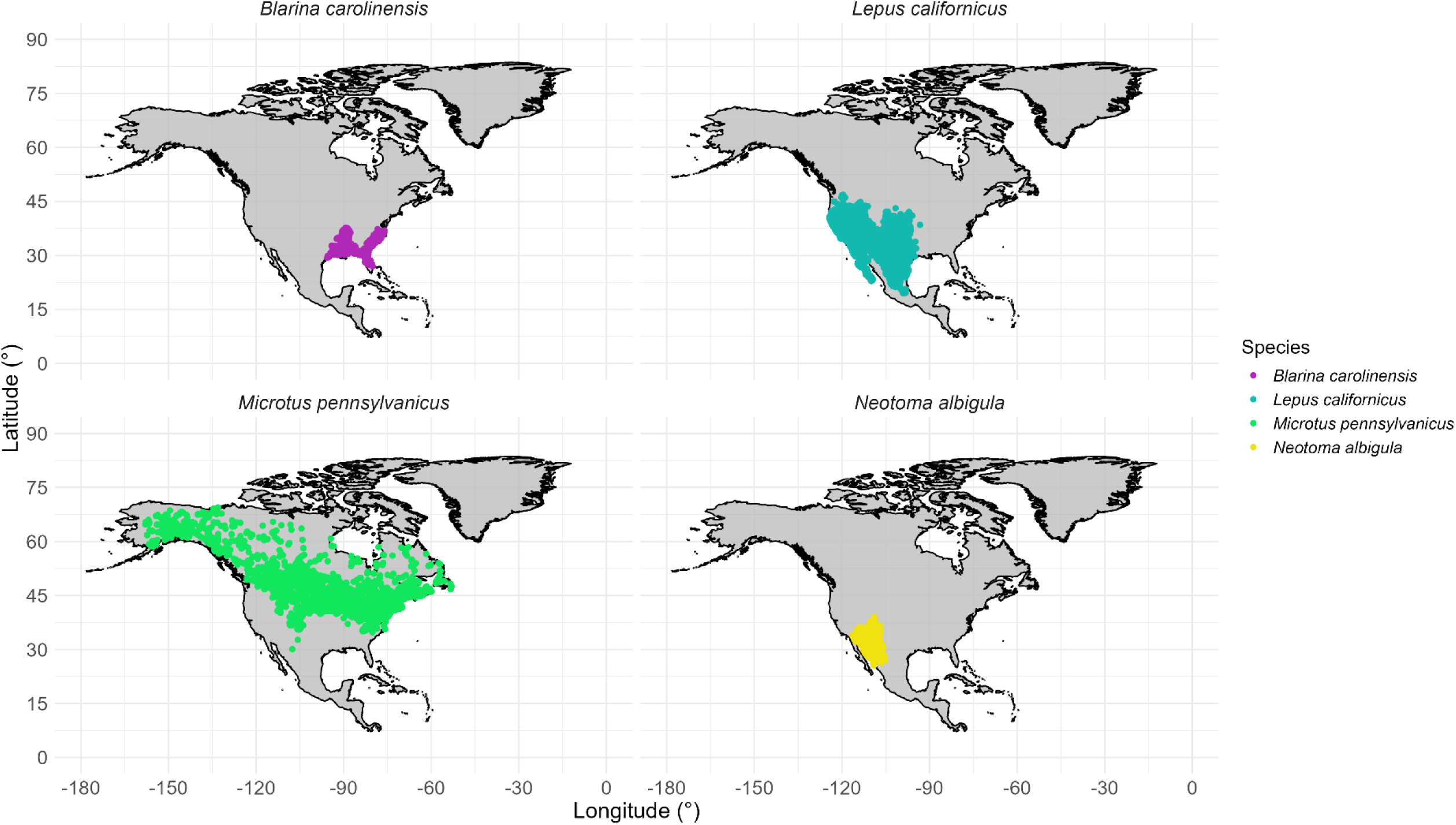
The distribution of GBIF occurrences for each species after excluding points based on their expert range map and spatial thinning. Map datum is WGS 1984.

**Figure S2.**
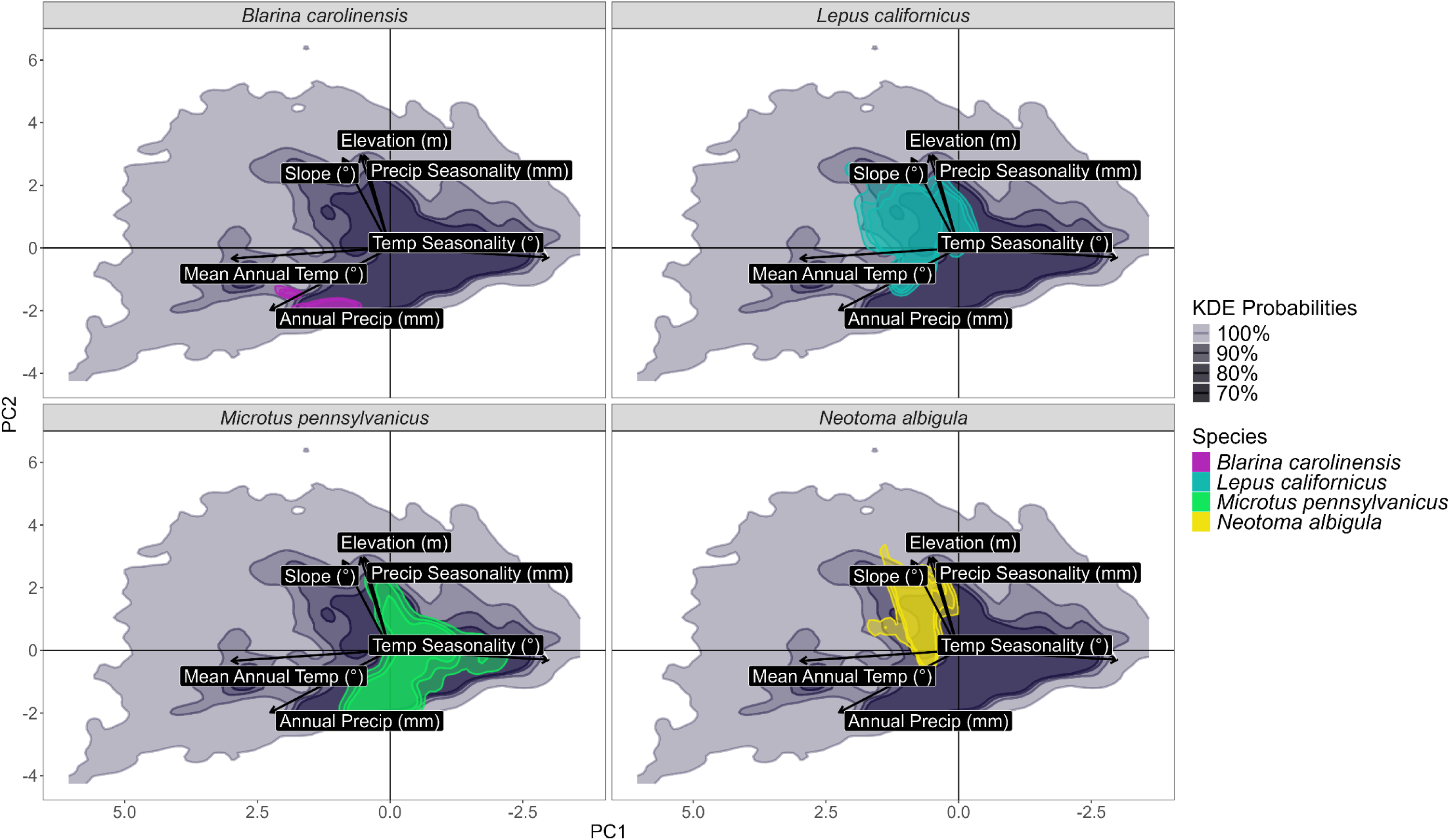
Present-day niche space of four real world species, obtained using GBIF occurrence records overlaid on the KDE of a random 99,999 point sample of North American environments (dark purple). The x-axis (PC1) scale has been reversed to align with^1^.

**Table S3.** Model evaluation and summary statistics for the ENM based on GBIF data for our four real species. Test AUC is averaged across the five-fold replicates. In addition, we show the three environmental variables with the highest permutation importance scores.

| Species | AUC | Permutational Importance (%) |  |  |
| --- | --- | --- | --- | --- |
|  |  | #1 | #2 | #3 |
| *Blarina carolinensis* | 0.77 | Annual Mean Temperature<br>(43.2) | Annual Precipitation<br>(42.7) | Elevation<br>(11.2) |
| *Lepus californicus* | 0.74 | Annual Mean Temperature<br>(60.5) | Temperature Seasonality<br>(17.6) | Annual Precipitation<br>(14.6) |
| *Microtus pennsylvanicus* | 0.70 | Annual Mean Temperature<br>(63.1) | Elevation<br>(14.9) | Temperature Seasonality<br>(11.4) |
| *Neotoma albigula* | 0.77 | Annual Mean Temperature<br>(65.8) | Temperature Seasonality<br>(17.5) | Elevation<br>(13.4) |

### SI.3 – Implementation (Extended)

Data manipulations were carried out with *dplyr* (version 1.1.2; Wickha4m et al. 2023*b*), *tidyr* (version 1.3.0; Wickham et al. 2023*a*) , and *stringr* (version 1.5.0; Wickham 2023). Geospatial data manipulations were performed with *terra* (version 1.7-78; Hijmans et al. 2022), *tidyterra* (version 0.6.0; Hernangómez 2023), and *sf* (version 1.0-16; Pebesma 2018). MaxEnt algorithm models were fitted using *dismo* (version 1.3-14; Hijmans et al. 2017), while niche comparisons were performed with *stats* (version 4.5.0; R Core Team 2025). Virtual species were created using *virtualspecies* (version 1.6; Leroy et al. 2016) with a modified version of *virtualspecies::generateSpFromFun* utilize *terra* (see code repository for details). Parallelization was implemented using *foreach* (version 1.5.2; Weston and Calaway 2015) and *doSNOW* (version 1.0.20; Calaway and Weston 2017). All base visualization were created with *ggplot2* (version 3.5.1; Wickham 2011), *ggdensity* (version 1.0.0; Otto and Kahle 2022), *tidyterra* and *viridis* (version 0.6.5). All analyses were conducted at the Ohio Supercomputer Center (1987) on the ‘Owens’ cluster.

### SI.4 – PCA model evaluation

Approximately half of our true PCAs showed a sharp drop off in variation explained after the first two principal components (Figure S3), indicating these models were well-represented by the first two axes, with a cumulative variance explained of 57.5% - 76.6% by these two axes. The remaining half all showed a drop off after the first three components.

**Figure S3.**
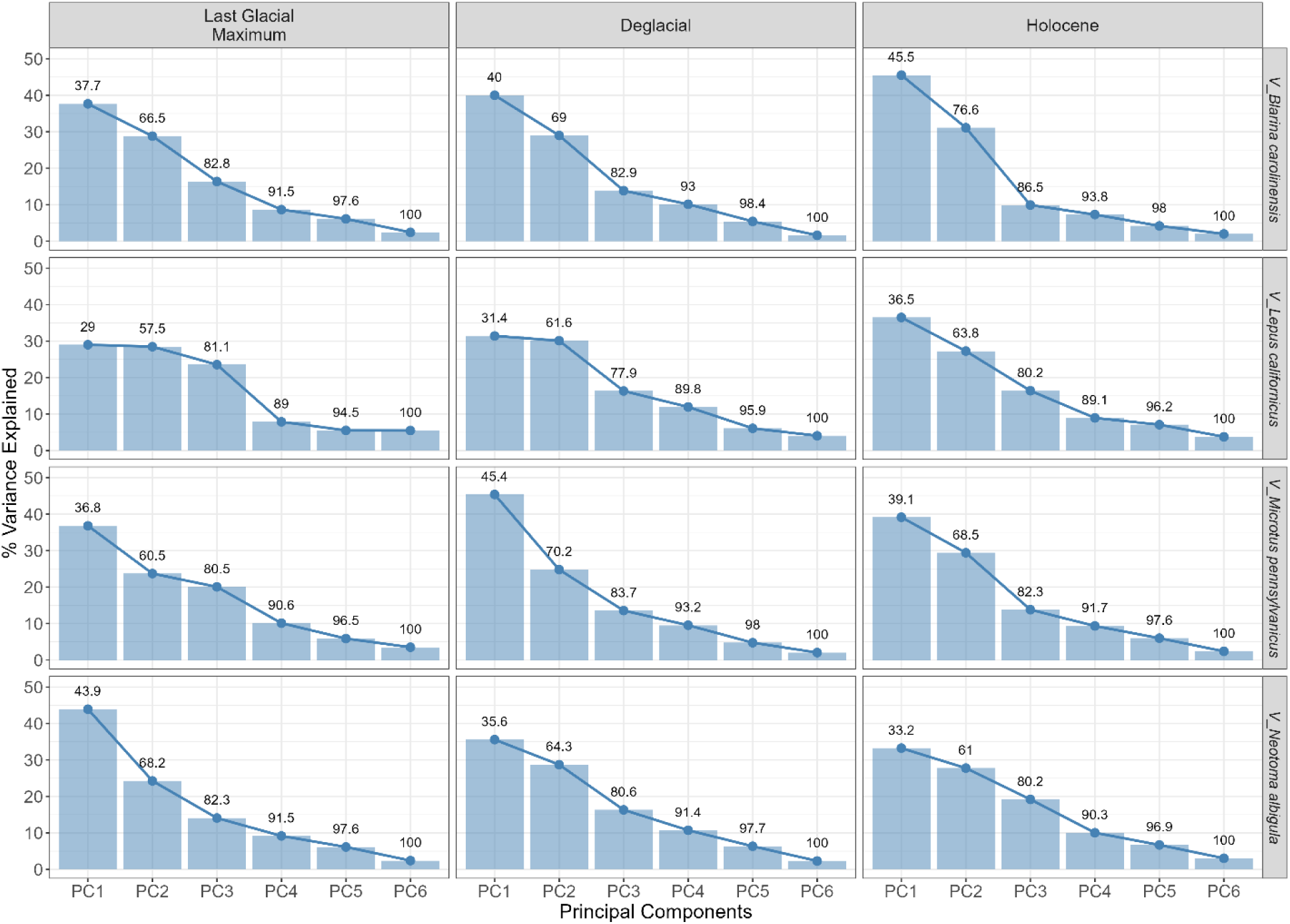
Variation explained by each principal component in the true Principal Component Analysis for each species and time period. Values above each bar show the cumulative variance explained by the principal components.

### SI.5 – Missing point impact assessment

Since the Last Glacial Maximum 23,000 years ago, glaciers have progressively retreated, accompanied by concurrent rises in sea level^16^; this has led to an expansion of inland habitats and a retreat of coastal habitats for small mammals in North America. As the observations for each species were sampled from the midpoint of each of the three periods, after temporal jittering, some locations may have been covered by glaciers or the sea. By masking out contemporaneous oceans, water bodies, and glaciers from our environmental rasters, temporal jittering can result in missing environmental data for some occurrences, which may impact model performance. To assess the impact of these missing points, we examined how NAs correlated with prediction error. The number of NAs was positively correlated with the level of temporal uncertainty, although the degree depended on both the midpoint and virtual species (Figure S4). NAs were positively correlated with prediction error during the LGM period for all virtual species.

Conversely, NAs appeared to have much less effect on Holocene prediction errors (Figure S4), even though this period had the replicate with the largest number of missing points (see *V_M. pennsylvanicus*). Excluding all replicates with any missing data points (Figure S5) showed very consistent trends between temporal uncertainty and prediction error to those seen when all data were included (main manuscript - Figure 3), albeit with *V_B. carolinensis* missing several levels of temporal uncertainty for both the LGM and the deglacial period.

**Figure S4.**
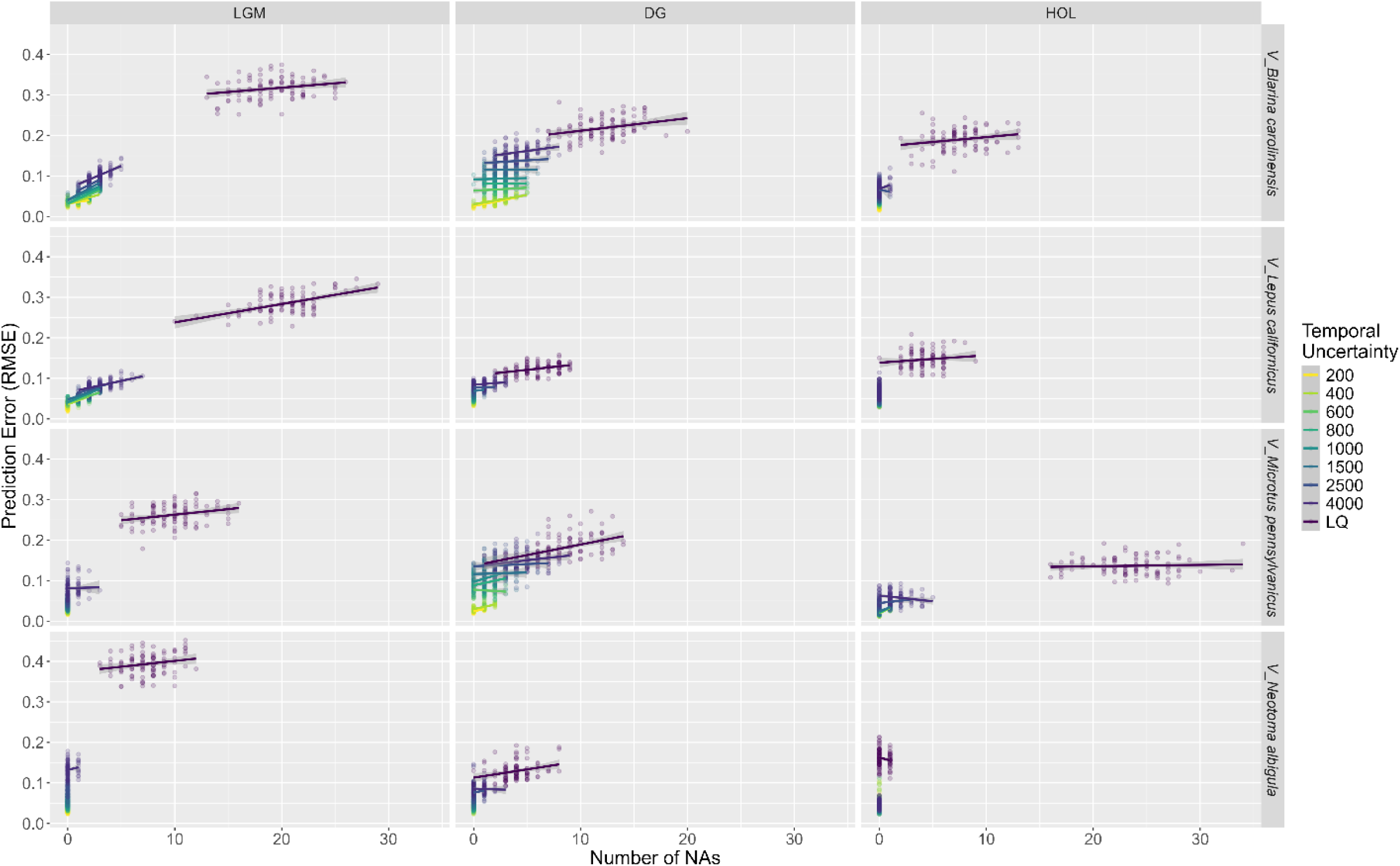
Prediction error plotted against the number of missing presence points for each species, time period and level of temporal uncertainty for each model replicate. Lines represent simple linear regressions (Gaussian errors) showing the relationship between prediction error and number of NAs.

**Figure S5.**
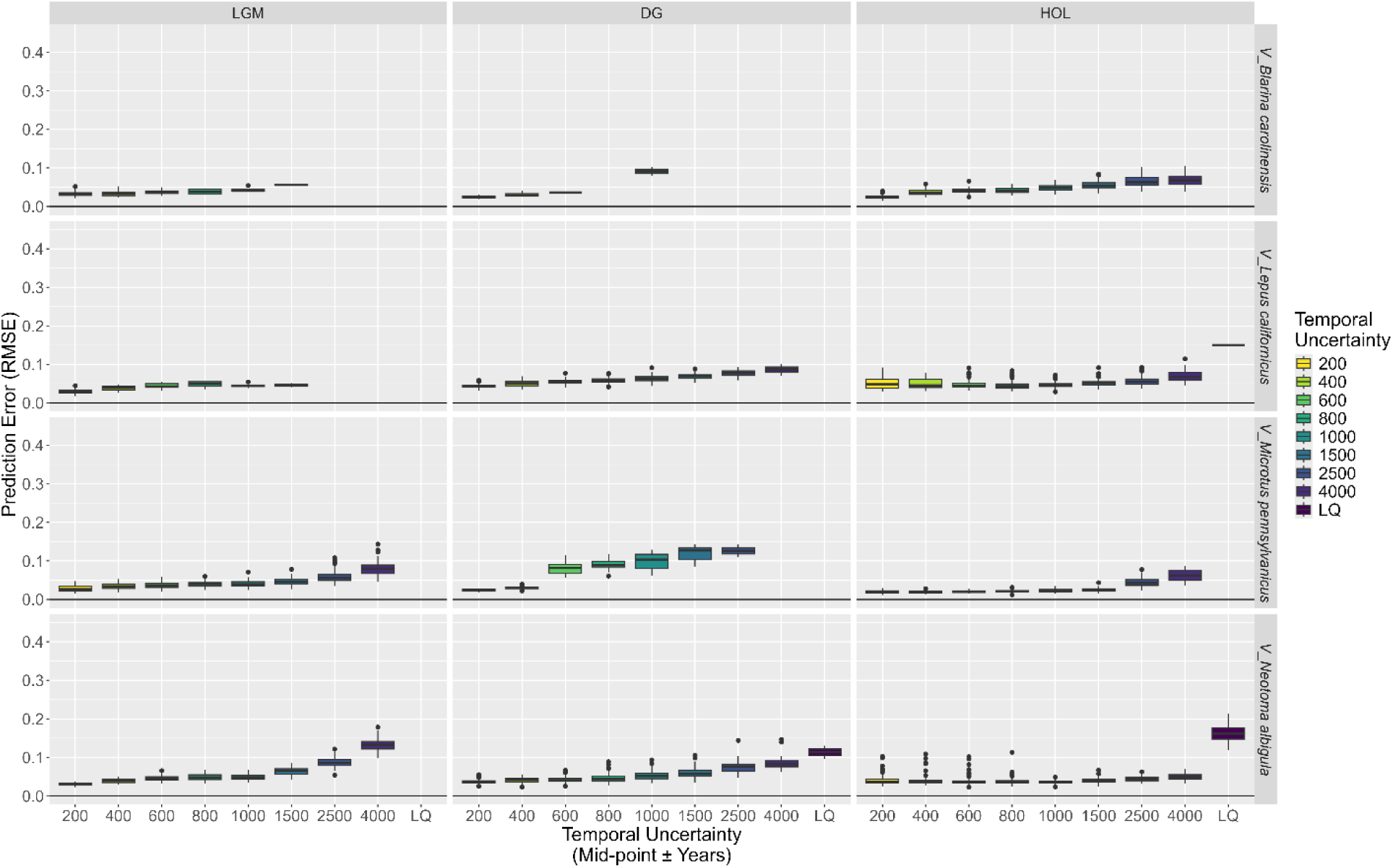
Average prediction error (root mean square error) for each virtual species and time period replicate, excluding any replicates that were missing data for any occurrences. The colour of the boxes indicating the temporal uncertainty range of the sample MaxEnt models.

Allowing replicates to have up to five missing values nearly perfectly reproduced the pattern seen in the full analysis, although four LQ temporal uncertainties were unrepresented (Figure S6). Allowing up to ten missing values reduced this to only two LQ temporal uncertainties unrepresented (*V_B. carolinensis* | LGM and *V_M. pennsylvanicus* | Holocene; Figure S7) – notably, the minimum number of NAs for these replicates was 13 and 16, respectively.

**Figure S6.**
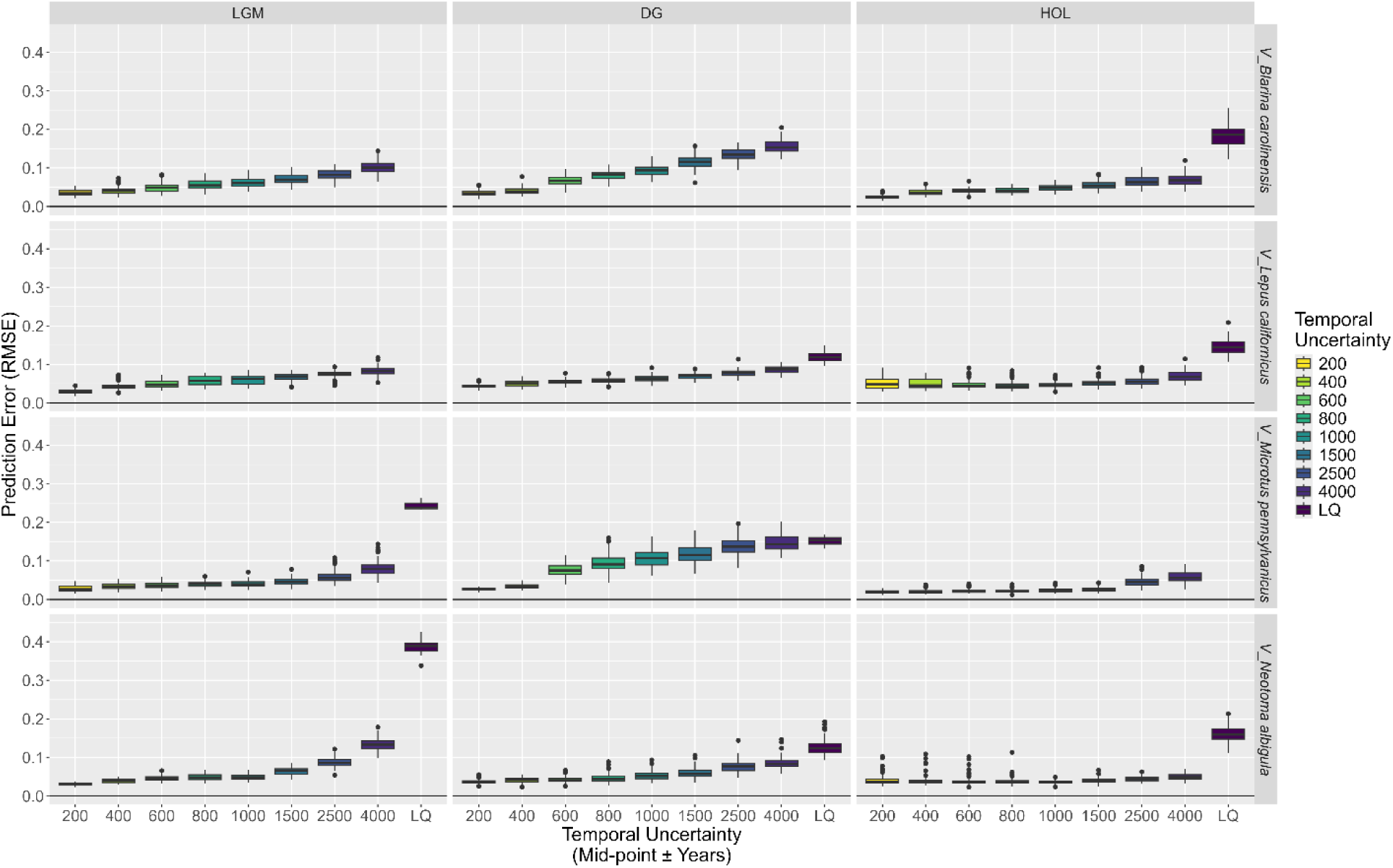
Average prediction error (root mean square error) for each virtual species and time period replicate, excluding replicates that were missing data for more than five occurrences. The colour of the boxes indicating the temporal uncertainty range of the sample MaxEnt models.

**Figure S7.**
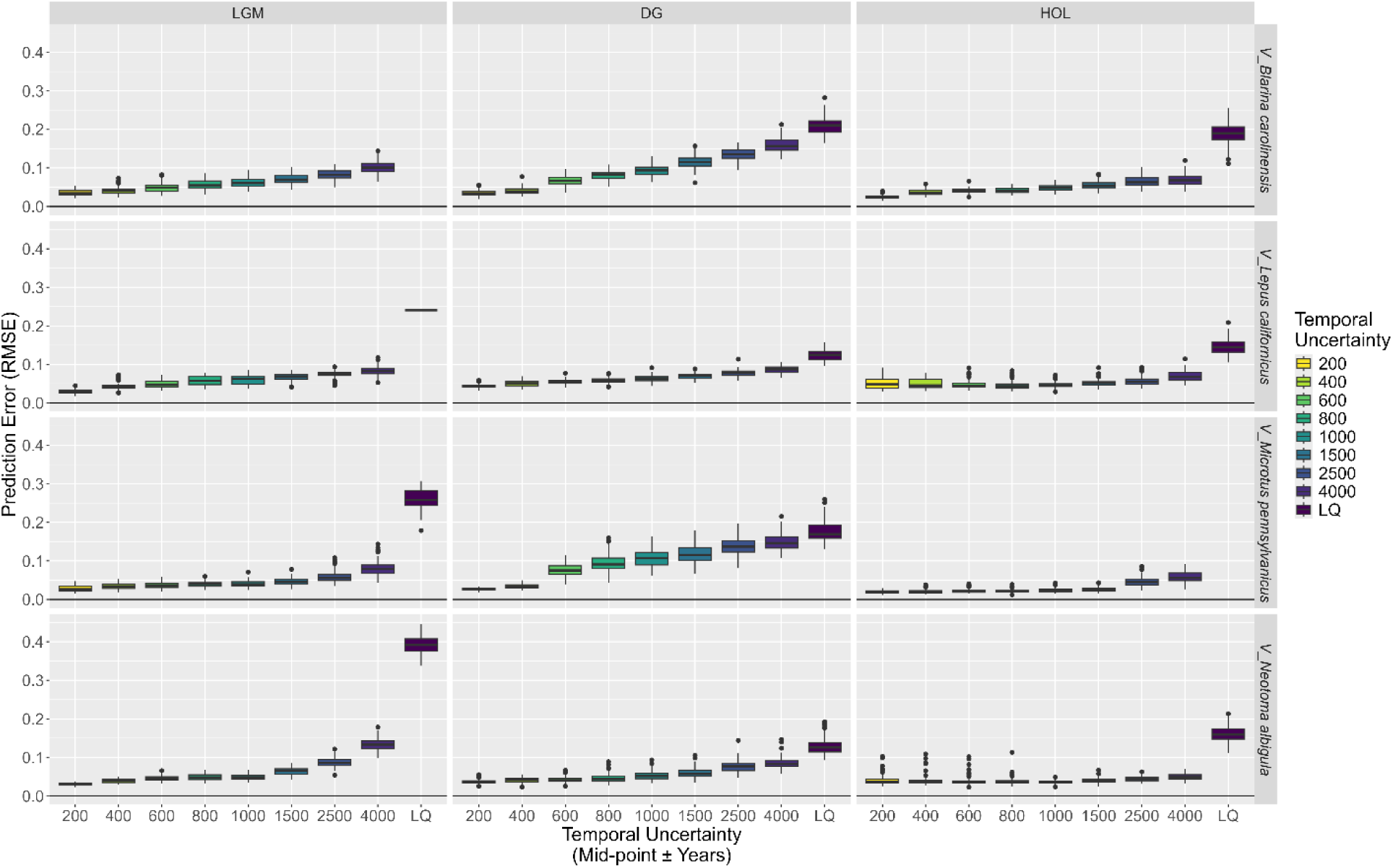
Average prediction error (root mean square error) for each virtual species and time period replicate, excluding replicates that were missing data for more than ten occurrences. The colour of the boxes indicates the temporal uncertainty range of the sample MaxEnt models.

The reproduction of the patterns observed in our main text, even when excluding large numbers of replicates, suggests that our conclusions remain valid. However, for *V_B. carolinensis* and *V_L. californicus* during the LGM in particular, NAs likely inflated prediction error when temporal uncertainty spanned the LQ, although the trends were in line with what *V_M. pennsylvanicus* and *V_N. albigula* showed when missing ≤ five data points. While it may be tempting to mask all environmental rasters down to a combined minimum area to ensure that no occurrences are lost, this would result in the loss of environments that were subject to the most pronounced environmental changes—which our study aimed to capture—and create artificial truncation of the virtual species niches. Conversely, expanding all environmental rasters to match the maximum possible extent could exaggerate the impact of temporal uncertainty, as extreme environments (e.g., large glacial expanses) would have been included in our model.

## References

Barbet-Massin, M., Jiguet, F., Albert, C.H. & Thuiller, W. (2012). Selecting pseudo-absences for species distribution models: how, where and how many? Methods in Ecology and Evolution, 3, 327–338.

Barido-Sottani, J., Van Tiel, N.M., Hopkins, M.J., Wright, D.F., Stadler, T. & Warnock, R.C. (2020). Ignoring fossil age uncertainty leads to inaccurate topology and divergence time estimates in time calibrated tree inference. Frontiers in Ecology and Evolution, 8, 183.

Barr, W.A. & Wood, B. (2024). Spatial sampling bias influences our understanding of early hominin evolution in eastern Africa. Nature Ecology & Evolution, 8, 2113–2120.

Beaumont, L.J., Graham, E., Duursma, D.E., Wilson, P.D., Cabrelli, A., Baumgartner, J.B., et al. (2016). Which species distribution models are more (or less) likely to project broad-scale, climate-induced shifts in species ranges? Ecological Modelling, 342, 135–146.

Behrensmeyer, A. & Chapman, R.E. (1993). Models and simulations of time-averaging in terrestrial vertebrate accumulations. Short Courses in Paleontology, 6, 125–149.

Behrensmeyer, A.K., Kidwell, S.M. & Gastaldo, R.A. (2000). Taphonomy and paleobiology. Paleobiology, 26, 103–147.

Bellvé, A.M., Syverson, V.J., Jarzyna, M.A. & Blois, J.L. (2025a). Preservation biases in the fossil record distort species ecological niche and distribution models. Ecography.

Bellvé, A.M., Wilmshurst, J.M., Wood, J.R., Whitehead, E., Scofield, R.P., Worthy, T.H., et al. (2025b). Burrowing into the past: Extending niche space models of procellariiform breeding grounds by merging fossil and historic data. Diversity and Distributions, 31, e70032.

Blaauw, M. (2012). Out of tune: the dangers of aligning proxy archives. Quaternary Science Reviews, 36, 38–49.

Blois, J.L., Bellvé, A.M., Jarzyna, M.A., Saupe, E.E. & Syverson, V.J. (2025). Paleobiogeographic insights gained from ecological niche models: progress and continued challenges. Paleobiology, 51, 8–28.

Bronk Ramsey, C. (2008). Radiocarbon Dating: Revolutions in understanding. Archaeometry, 50, 249–275.

Canteri, E., Brown, S.C., Schmidt, N.M., Heller, R., Nogues-Bravo, D. & Fordham, D.A. (2022). Spatiotemporal influences of climate and humans on muskox range dynamics over multiple millennia. Global Change Biology, 28, 6602–6617.

Chiarenza, A.A., Mannion, P.D., Lunt, D.J., Farnsworth, A., Jones, L.A., Kelland, S.-J., et al. (2019). Ecological niche modelling does not support climatically-driven dinosaur diversity decline before the Cretaceous/Paleogene mass extinction. Nat Commun, 10, 1091.

Clark, P.U., Shakun, J.D., Baker, P.A., Bartlein, P.J., Brewer, S., Brook, E., et al. (2012). Global climate evolution during the last deglaciation. Proceedings Of The National Academy Of Sciences Of The United States Of America, 109, E1134–42.

Darroch, S.A.F., Fraser, D. & Casey, M.M. (2021). The preservation potential of terrestrial biogeographic patterns. Proceedings of the Royal Society B: Biological Sciences, 288, 20202927.

Davis, E.B., McGuire, J.L. & Orcutt, J.D. (2014). Ecological niche models of mammalian glacial refugia show consistent bias. Ecography, 37, 1133–1138.

Dietl, G.P. & Flessa, K.W. (2011). Conservation paleobiology: putting the dead to work. Trends in Ecology & Evolution, 26, 30–37.

Dos Reis, M., Thawornwattana, Y., Angelis, K., Telford, M.J., Donoghue, P.C. & Yang, Z. (2015). Uncertainty in the timing of origin of animals and the limits of precision in molecular timescales. Current biology, 25, 2939–2950.

Drummond, A.J. & Stadler, T. (2016). Bayesian phylogenetic estimation of fossil ages. Philosophical Transactions of the Royal Society B: Biological Sciences, 371, 20150129.

Fielding, A.H. & Bell, J.F. (1997). A review of methods for the assessment of prediction errors in conservation presence/absence models. Environmental conservation, 24, 38–49.

Flannery-Sutherland, J.T., Elsler, A., Farnsworth, A., Lunt, D.J. & Benton, M.J. (2025). Landscape-explicit phylogeography illuminates the ecographic radiation of early archosauromorph reptiles. Nat Ecol Evol, 9, 1138–1152.

Foffa, D., Dunne, E.M., Chiarenza, A.A., Wynd, B.M., Farnsworth, A., Lunt, D.J., et al. (2025). Climate drivers and palaeobiogeography of lagerpetids and early pterosaurs. Nat Ecol Evol, 9, 1359–1372.

Fordham, D.A., Brown, S.C., Akçakaya, H.R., Brook, B.W., Haythorne, S., Manica, A., et al. (2022). Process-explicit models reveal pathway to extinction for woolly mammoth using pattern-oriented validation. Ecology Letters, 25, 125–137.

Fordham, D.A., Jackson, S.T., Brown, S.C., Huntley, B., Brook, B.W., Dahl-Jensen, D., et al. (2020). Using paleo-archives to safeguard biodiversity under climate change. Science, 369, eabc5654.

Graham, R.W. (1993). Processes of time-averaging in the terrestrial vertebrate record. Short Courses in Paleontology, 6, 102–124.

Grether, G.F., Finneran, A.E. & Drury, J.P. (2024). Niche differentiation, reproductive interference, and range expansion. Ecology Letters, 27, e14350.

Herrando-Pérez, S. & Stafford Jr., T.W. (2025). Making vertebrate fossil radiocarbon dates more useful for global scientific research. Journal of Quaternary Science, 40, 1309–1335.

Herrando-Pérez, S. & Stafford, T.W. (2025). Making vertebrate fossil radiocarbon dates more useful for global scientific research. J Quaternary Science, 40, 1309–1335.

Hoban, S., Dawson, A., Robinson, J.D., Smith, A.B. & Strand, A.E. (2019). Inference of biogeographic history by formally integrating distinct lines of evidence: genetic, environmental niche and fossil. Ecography, 42, 1991–2011.

Inman, R., Franklin, J., Esque, T. & Nussear, K. (2018). Spatial sampling bias in the Neotoma paleoecological archives affects species paleo-distribution models. Quaternary Science Reviews, 198, 115–125.

Johnson, C.N., Balmford, A., Brook, B.W., Buettel, J.C., Galetti, M., Guangchun, L., et al. (2017). Biodiversity losses and conservation responses in the Anthropocene. Science, 356, 270–275.

Johnson, T., Beckerman, A., Childs, D., Webb, T., Evans, K., Griffiths, C., et al. (2024). Revealing uncertainty in the status of biodiversity change. Nature, 628, 788–794.

Karger, D.N., Nobis, M.P., Normand, S., Graham, C.H. & Zimmermann, N.E. (2023). CHELSA-TraCE21k – high-resolution (1 km) downscaled transient temperature and precipitation data since the Last Glacial Maximum. Climate of the Past, 19, 439–456.

Leroy, B., Meynard, C.N., Bellard, C. & Courchamp, F. (2016). virtualspecies, an R package to generate virtual species distributions. Ecography, 39, 599–607.

Lima-Ribeiro, M.S., Moreno, A.K.M., Terribile, L.C., Caten, C.T., Loyola, R., Rangel, T.F., et al. (2017). Fossil record improves biodiversity risk assessment under future climate change scenarios. Diversity and Distributions, 23, 922–933.

Maguire, K.C., Nieto-Lugilde, D., Blois, J.L., Fitzpatrick, M.C., Williams, J.W., Ferrier, S., et al. (2016). Controlled comparison of species- and community-level models across novel climates and communities. Proceedings of the Royal Society B: Biological Sciences, 283, 20152817.

Maguire, K.C., Nieto-Lugilde, D., Fitzpatrick, M.C., Williams, J.W. & Blois, J.L. (2015). Modeling species and community responses to past, present, and future episodes of climatic and ecological change. Annual Review of Ecology, Evolution, and Systematics, 46, 343–368.

Maxwell, S.J., Hopley, P.J., Upchurch, P. & Soligo, C. (2018). Sporadic sampling, not climatic forcing, drives observed early hominin diversity. Proceedings of the National Academy of Sciences, 115, 4891–4896.

Meersch, V.V. der, Armstrong, E., Mouillot, F., Duputié, A., Davi, H., Saltré, F., et al. (2025). Paleorecords Reveal Biological Mechanisms Crucial for Reliable Species Range Shift Projections Amid Rapid Climate Change. Ecology Letters, 28, e70080.

Meltzer, D.J. & Mead, J.I. (1985). Dating late Pleistocene extinctions: theoretical issues, analytical bias, and substantive results. Environments and extinctions: Man in late glacial North America, 145–73.

Myers, C.E., Stigall, A.L. & Lieberman, B.S. (2015). PaleoENM: applying ecological niche modeling to the fossil record. Paleobiology, 41, 226–244.

Naughtin, S.R., Castilla, A.R., Smith, A.B., Strand, A.E., Dawson, A., Hoban, S., et al. (2025). Integrating genomic data and simulations to evaluate alternative species distribution models and improve predictions of glacial refugia and future responses to climate change. Ecography, 2025, e07196.

Noto, C.R. (2011). Hierarchical control of terrestrial vertebrate taphonomy over space and time: discussion of mechanisms and implications for vertebrate paleobiology. In: Taphonomy: Process and bias through time, Aims & Scope Topics in Geobiology Book Series. Springer, Dordrecht, pp. 287–336.

O’keefe, F.R., Dunn, R.E., Weitzel, E.M., Waters, M.R., Martinez, L.N., Binder, W.J., et al. (2023). Pre–Younger Dryas megafaunal extirpation at Rancho La Brea linked to fire-driven state shift. Science, 381, eabo3594.

Parker, A.K., McHorse, B.K. & Pierce, S.E. (2018). Niche modeling reveals lack of broad-scale habitat partitioning in extinct horses of North America. Palaeogeography, palaeoclimatology, palaeoecology, 511, 103–118.

Parnell, A.C., Buck, C.E. & Doan, T.K. (2011). A review of statistical chronology models for high-resolution, proxy-based Holocene palaeoenvironmental reconstruction. Quaternary Science Reviews, 30, 2948–2960.

Payne, A.R.D., Mannion, P.D., Lloyd, G.T. & Davis, K.E. (2024). Decoupling speciation and extinction reveals both abiotic and biotic drivers shaped 250 million years of diversity in crocodile-line archosaurs. Nature Ecology & Evolution, 8, 121–132.

Phelps, L.N., Broennimann, O., Manning, K., Timpson, A., Jousse, H., Mariethoz, G., et al. (2020). Reconstructing the climatic niche breadth of land use for animal production during the African Holocene. Global Ecology and Biogeography, 29, 127–147.

Pol, D. & Norell, M.A. (2006). Uncertainty in the age of fossils and the stratigraphic fit to phylogenies. Systematic Biology, 55, 512–521.

R Core Team. (2025). R: A language and environment for statistical computing.

Raiter, K.G. & Hawlena, D. (2024). Managing multiple uncertainties in species distribution modelling. Diversity and Distributions, 30, e13857.

Razgour, O., Juste, J., Ibáñez, C., Kiefer, A., Rebelo, H., Puechmaille, S.J., et al. (2013). The shaping of genetic variation in edge-of-range populations under past and future climate change. Ecology Letters, 16, 1258–1266.

Reimer, P.J., Austin, W.E.N., Bard, E., Bayliss, A., Blackwell, P.G., Bronk Ramsey, C., et al. (2020). The IntCal20 Northern Hemisphere Radiocarbon Age Calibration Curve (0–55 cal kBP). Radiocarbon, 62, 725–757.

Reside, A.E., VanDerWal, J.J., Kutt, A.S. & Perkins, G.C. (2010). Weather, Not Climate, Defines Distributions of Vagile Bird Species. PLOS ONE, 5, e13569.

Rindel, D.D., Moscardi, B.F. & Perez, S.I. (2021). The distribution of the guanaco (Lama guanicoe) in Patagonia during Late Pleistocene–Holocene and its importance for prehistoric human diet. The Holocene, 31, 644–657.

Rstudio Team. (2025). RStudio: integrated development for R.

Saltré, F., Brook, B.W., Rodríguez-Rey, M., Cooper, A., Johnson, C.N., Turney, C.S., et al. (2015). Uncertainties in dating constrain model choice for inferring extinction time from fossil records. Quaternary Science Reviews, 112, 128–137.

Stigall, A.L. (2012). Using ecological niche modelling to evaluate niche stability in deep time. Journal of Biogeography, 39, 772–781.

Syverson, V.J.P., Goring, S.J., Cullen, N., Jarzyna, M.A., Bellvé, A.M., Martindale, A., et al. (2026). Updated chronologies for North American small mammal fossil localities in the Neotoma Paleoecology Database. Sci Data.

Timmermann, A., Yun, K.-S., Raia, P., Ruan, J., Mondanaro, A., Zeller, E., et al. (2022). Climate effects on archaic human habitats and species successions. Nature, 604, 495–501.

Tomašových, A., Kowalewski, M., Nawrot, R., Scarponi, D. & Zuschin, M. (2024). Abundance–diversity relationship as a unique signature of temporal scaling in the fossil record. Ecology Letters, 27, e14470.

Veloz, S.D., Williams, J.W., Blois, J.L., He, F., Otto-Bliesner, B. & Liu, Z. (2012). No-analog climates and shifting realized niches during the late quaternary: implications for 21st-century predictions by species distribution models. Global Change Biology, 18, 1698–1713.

Walker, M. (2013). Quaternary dating methods. John Wiley & Sons.

Walls, B.J. & Stigall, A.L. (2011). Analyzing niche stability and biogeography of Late Ordovician brachiopod species using ecological niche modeling. Palaeogeography, Palaeoclimatology, Palaeoecology, 299, 15–29.

Waterson, A.M., Schmidt, D.N., Valdes, P.J., Holroyd, P.A., Nicholson, D.B., Farnsworth, A., et al. (2016). Modelling the climatic niche of turtles: a deep-time perspective. Proceedings of the Royal Society B: Biological Sciences, 283, 20161408.

Wendt, J.A.F., McWethy, D.B., Widga, C. & Shuman, B.N. (2022). Large-scale climatic drivers of bison distribution and abundance in North America since the Last Glacial Maximum. Quaternary Science Reviews, 284, 107472.

Williams, J.W., Kharouba, H.M., Veloz, S., Vellend, M., McLachlan, J., Liu, Z., et al. (2013). The ice age ecologist: testing methods for reserve prioritization during the last global warming. Global Ecology and Biogeography, 22, 289–301.

Wright, D.K. (2017). Accuracy vs. Precision: Understanding Potential Errors from Radiocarbon Dating on African Landscapes. African Archaeological Review, 34, 303–319.

Zurell, D., Berger, U., Cabral, J.S., Jeltsch, F., Meynard, C.N., Münkemüller, T., et al. (2010). The virtual ecologist approach: simulating data and observers. Oikos, 119, 622–635.

